# A mathematical model relates intracellular TLR4 oscillations to sepsis progression

**DOI:** 10.1101/164137

**Authors:** Razvan C. Stan, Francisco G. Soriano, Maristela M. de Camargo

**Affiliations:** Laboratory of Molecular Immunoregulation, Institute of Biomedical Sciences, University of São Paulo, CEP 05508-900, São Paulo, Brazil; University Hospital, University of Sao Paulo, CEP 05508-000, São Paulo, Brazil; School of Medicine, University of Sao Paulo, CEP 01246-904, São Paulo, Brazil

**Keywords:** Gram negative infections, inflammation, intracellular trafficking

## Abstract

Oscillations drive many biological processes and their modulation is determinant for various pathologies. In sepsis syndrome, Toll-like receptor 4 (TLR4) is a key sensor for signaling the presence of Gram-negative bacteria. Its expression and activity, along with its intracellular trafficking rates shift the equilibrium between the pro‐ and anti-inflammatory downstream signaling cascades, leading to either the physiological resolution of the bacterial stimulation or to sepsis. We hypothesize that the initial *tlr4* expression in patients diagnosed with sepsis and TLR4 dynamic concentration changes on the cell membrane or intracellularly, dictates how the sepsis syndrome is initiated. Using a set of three differential equations, we defined the TLR4 flux between relevant cell organelles. We obtained three different regions in the phase space: 1. a limit-cycle describing unstimulated physiological oscillations, 2. a fixed-point attractor resulting from moderate LPS stimulation that is resolved and 3. a double-attractor resulting from sustained LPS stimulation that leads to sepsis. We tested the models against hospital data of sepsis patients and we correctly evaluate the clinical outcome of these patients.

## 1. Introduction

The immune system is replete with oscillations of various parameters needed for mounting an appropriate response upon stimulation. These include periodic variations in cytokine concentrations following antigen challenge [1], oscillations in the concentrations of Ca^2+^ or reactive oxygen species in neutrophils [2], oscillations in nuclear factor kB activity following stimulation by tumor necrosis factor alpha [3] etc. Importantly, the frequency and amplitude of these oscillations vary with inflammatory status and may have diagnostic value [4]. We have sought to determine whether such periodic oscillations are also manifested in the intracellular expression and trafficking of key pathogen sensors. This situation is particularly relevant due to cyclical and cross-inhibiting pro‐ and anti-inflammatory responses these sensors elicit upon stimulation. We have herein focused on the sepsis syndrome, a life-threatening clinical disorder that encompasses the physiological reactions to invading pathogens and/or their toxins, and that is responsible for high mortality rates [5]. TLR4 is a key recognition receptor for Gram-negative bacteria, and together with other members of the Toll-family, serves as a link between innate and adaptive immunity [6]. The early involvement of surface TLR4 in mediating the systemic responses to both invading pathogens and endogenous ligands is essential for sepsis pathogenesis [7], and as such it may serve as a crucial initial sepsis biomarker. In particular, TLR4 experiences a significant upregulation in mRNA production and presentation to the cell surface at the initial stages of sepsis in both humans and experimental models [8]. It is not evident however whether such increase positively correlates with the later progression into septic shock, as patients show similar TLR4 protein levels when compared to less severe septic stages [9,10]. Similarly, in experimental models of endotoxin tolerance, while the TLR4 concentrations on the surface of human peripheral blood mononuclear cells remain unchanged, the overall responsiveness to secondary LPS stimulation decreases [11]. Such effects may however be related to the disproportionate modulation of the distinct inflammatory signaling branches upon TLR4 activation. Throughout the continuum of sepsis, complete TLR4 signaling includes not only the initial surface-bound pro-inflammatory signaling, but also its subsequent endocytosis and intracellular trafficking. This results in competing endosomal anti-inflammatory cytokine production and further in either receptor recycling to cell membrane or signal termination within endolysosomes [12]. Initial responsiveness to LPS is therefore regulated by the concentrations of cell surface TLR4 that depend in turn on both TLR4 trafficking from the Golgi apparatus to the plasma membrane and on the amount of TLR4 internalized into endosomes [13-15].

## 2. Model construction

An emerging theme in TLR4 signaling posits that its cellular localization is determinant for its functions [16-18]. An overview of the known TLR4 intracellular trafficking routes that influence its signaling is presented in Figure 1.

**Fig. 1.**
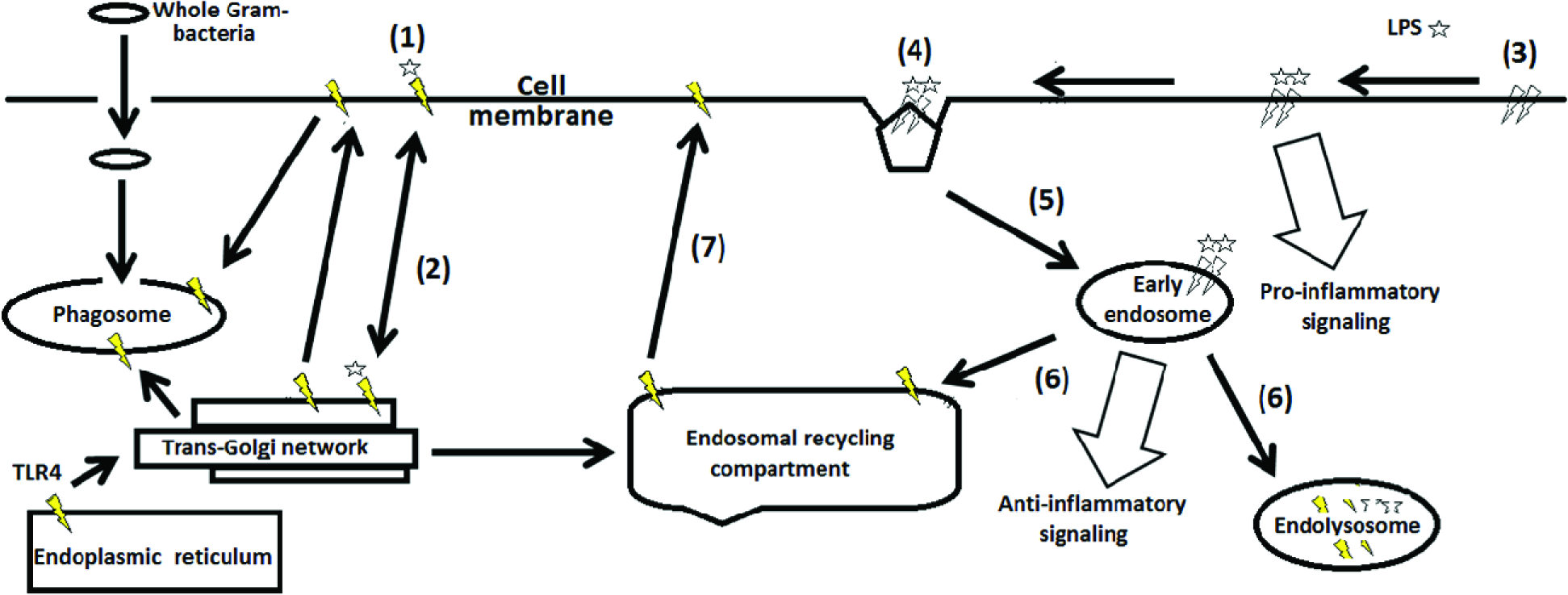
General scheme for TLR4 distribution and activation between different cellular compartments. Star symbols represent single LPS ligands; yellow thunder symbols depict single TLR4, white thunders describe signaling-competent TLR4 dimers. Numbers indicate the steps of LPS binding, followed by TLR4 trafficking and signaling events. and are described in text. For clarity, multiple TLR4 modulators present on the cell surface or intracellularly are not shown.

Upon endotoxin stimulation, initial TLR4 immobilization (step 1) may lead to monomeric LPS being internalized and trafficked to the Golgi apparatus within seconds of stimulation, without activating TLR4. This process serves mainly to limit the impinging endotoxin pool that may induce TLR4 hyperactivation [19]. This is followed by TLR4 clustering (step 3) with monomeric LPS [20]. Internalization by either clathrin‐ and dynamin-mediated processes [21] results in a switch in TLR4 signaling pathways by means of different adaptors (step 4). In a first signaling wave occurring at the cell surface, TIRAP (toll-interleukin 1 receptor (TIR) domain containing adaptor protein)-MyD88 (myeloid differentiation primary response gene (88)-assisted pro-inflammatory cytokine production is initiated. Provided that TLR4 endocytosis has occurred, signaling continues with TRAM (TRIF-related adaptor molecule)-TRIF (TIR-domain-containing adapter-inducing interferon-β) adaptor complex formation and subsequent induction of type-I interferons [22] from early endosomes (step 5). From the early endosomes, for the signal to be terminated (step 6), the TLR4 complex is ubiquitinated, marked for lysosomal degradation and loaded with associated antigens for the presentation to CD4+ T cells [23]. Alternatively, TLR4 can be recycled for new signaling cycles back to the cell surface via the endosomal recycling compartment (step 7). It is important to note that this scheme does not include phagosome signaling of whole Gram negative bacteria, nor the additional TLR4 subpopulation that trafficks from ERC to phagosome after LPS stimulation. TLR4 expression and cell surface presentation is crucial for initiating the bacterial presence signaling cascade, such that blocking surface TLR4 affords protection from induced infections that are otherwise lethal [24], whereas experimentally increased TLR4 expression through gene dosage exacerbate the pro-inflammatory signaling [25], as does inhibition of TRL4 endocytosis and its endosomal sorting [23]. In the absence of LPS stimulation, steady state concentrations for *tlr4* mRNA in human monocytes and macrophages oscillate within a factor of 3 from initial values (days 1-5) [26]. Under limit-cycle unstimulated physiological oscillations, cell surface TLR4 are present in low concentrations in macrophages or are undetectable in dendritic cells, with most resident TLR4 being distributed in the Golgi apparatus [27]. Rapid TLR4 mobilization to cell membrane follows LPS activation [28], canceling the downregulation present in physiological conditions that serves to desensitize cells to low endotoxin levels. While the overall sequence of TLR4 activation has been elucidated, the rates of TLR4 trafficking are not quantified, nor are available absolute numbers for TLR4 expression on cell surfaces.

In order to simulate in silico the initial TLR4 trafficking events between the endosomal recycling compartment (ERC) and the Trans-Golgi network (TGN) to and from cell surface and within the early endosomes-endolysosome (EE) system, we have constructed a dynamic model based on the three ordinary differential equations presented below:

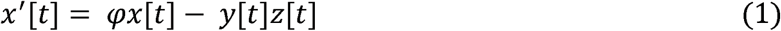

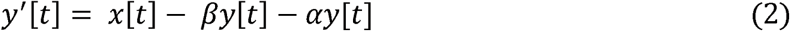

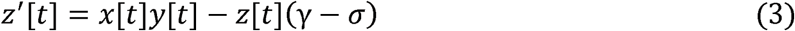

where: x = concentration of TLR4 in TGN and ERC, y = concentration of TLR4 in endosomes/endolysosomes (EE), z = concentration of TLR4 on cell surface, □ = rate of TLR4 mRNA production, β = rate of TLR4 trafficked to lysosomes from endosomes, α = rate of TLR4 retroactively trafficked to ERC from endosomes, γ = rate of TLR4 on cell surface trafficked to TGN, σ = rate of TLR4 on cell surface trafficked to endosomal system.

The TLR4 flux in the system as indicated by equation (1) is influenced by the TRAM distribution within ERC that shifts onto the enlarged CD14/LPS-positive endosomes upon TLR4 activation [18]. The adaptor TRAM is also constitutively present at the plasma membrane anchored at a N-terminal myristoylation site and traffics concomitantly the TLR4 signaling complex unidirectionaly to the endosomal system [29]. This synergy allows for the antiinflammatory signaling phase to take preponderance, possibly due to unique TLR4 conformation brought on by the endosomal acidic environment, as previously proposed [30]. These events are dominant after about 30 minutes upon LPS stimulation [21], allowing for TLR4 to traffic in a first stage mostly bidirectionally from the ERC to EE (equation (2). The small GTPase Rab7b is a key regulator of TLR4 intracellular trafficking that is upregulated upon LPS exposure in the early endosomes leading to its transport to either late endosomes/lysosomes for signal termination or to ERC [31], as represented by equation (3). In Rab7b-silenced macrophages, after LPS stimulation, continued TLR4 presence only in the EE system has adverse effects as to its prolonged anti-inflammatory signaling [32]. Equation (3) describes the TLR4 cell surface concentration changes as the difference between the pool of available TLR4 in TGN + ERC and in EE, and the TLR4 that is actively being prevented from clustering on the cell surface (so as to increase downstream signaling), be it directly from the surface towards TGN (parameter γ) or towards EE (parameter σ).

As mentioned, in unstimulated cells, TLR4 mRNA expression is not statistically different across days 1-5 in human macrophages [26]. Furthermore, in a shock serum model, spleen TLR4 mRNA expression did not show significant daily fluctuations. [33] In the absence of available data from the literature, the parameter values for □, β, γ σ and □ were varied until a stable limit cycle was attained, corresponding to physiological fluctuations in TLR4 expression. The first 4 parameters were kept constant to reflect the steady-state, non-stimulated oscillations in the TLR4 intracellular trafficking, while the □-parameter that has been determined experimentally [16] was allowed to vary.

## 3. Results

Using equations 1-3, we sought to model the cellular regimes that are impacted by the overall TLR4 sensitivity to LPS, as reflected by the initial rate of *tlr4* mRNA synthesis, upon sepsis diagnosis and prior to clinical intervention. The □-parameter augments markedly in experimental models of sepsis and directly correlates with mortality, with peak increases between 1-3 hours post sepsis induction [8]. We defined three regions in the phase space for the plasma membrane and intracellular TLR4 distribution, based on the variations of □-parameter drawn from Supplemental Table 2: (i) a steady-state with TLR4 expression and concentration oscillating within a narrow margin throughout the relevant cell compartments, (ii) a low to medium *tlr4* mRNA production following LPS stimulation that results in an initial increase of TLR4 concentration on the cell surface and subsequently in the endosomal system, followed by a regulated decrease, (iii) a third, high *tlr4* mRNA output matching increasing LPS stimulation where TLR4 concentrations oscillate stably and irreversibly on the cell surface and within the EE. The variations in *tlr4* mRNA measured in the patients served as the initial parameter (□) to be changed, responsible for initial TLR4 distribution within the relevant cell compartments. TLR4 is unique among other pathogen-recognition receptors in that its intracellular trafficking is determinant for the inflammatory signaling it initiates. As such, oscillations in its concentration within various relevant cell compartments will dictate the timing and preponderance of the pro‐ and anti-inflammatory responses. Depending on initial conditions and rate changes, the ensuing orbits either approach stable fixed points or undergo variations, each having a different physiological interpretation, as presented in Figure 2.

**Fig. 2.**
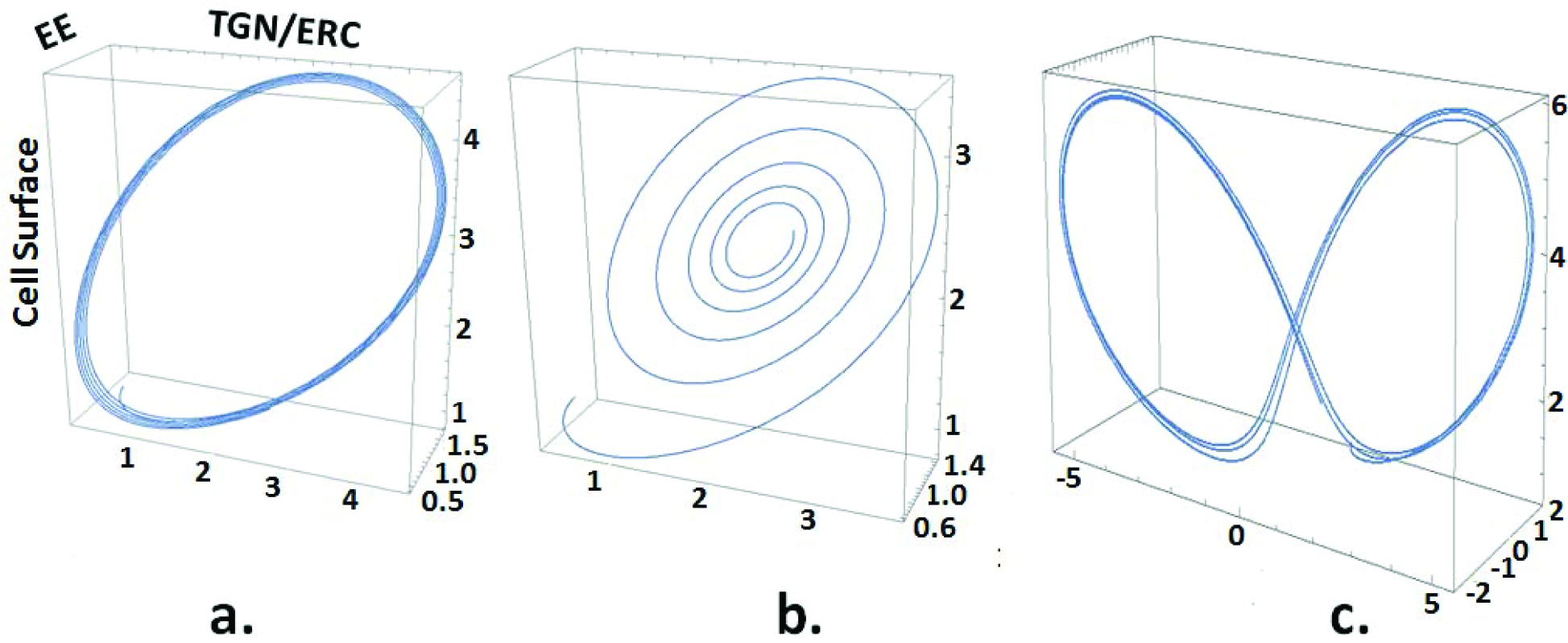
Simulated TLR4 cellular distribution during sepsis. a. Attractive limit cycle representing steady-state oscillations. b. Fixed-point attractor obtained following a low to medium (□ < 1.2) *tlr4* mRNA increase that temporarily augments TLR4 concentrations on the cell surface and thereafter within the EE system. c. Double-attractor obtained upon increasing *tlr4* mRNA, that leads to high TLR4 concentrations oscillating indeterminately between EE and cell membrane. X axis = TLR4 concentration in TGN/ERC. Y axis = concentration of TLR4 in EE. Z axis = concentration of TLR4 on cell surface.

a. Physiological variations in TLR4 concentrations In all simulations, we assumed that initial expression levels on the cell surface, TGN/ERC and EE are similarly low. For steady-state conditions, we proposed that TLR4 concentration oscillations are of low amplitude, reflecting the experimental data on *tlr4* mRNA in human monocytes in vitro (25). A stable limit cycle is achieved with □ = 1.2, β = 3.6, α = 1.2, γ = 2.4, σ = 1.3 (Fig 2, panel a).
b. Sepsis progression and resolution We surmised that following a moderate LPS stimulation, TLR4 levels initially increase in order to proportionally signal the Gram-negative bacterial presence, as previously documented in septic human patients [8]. A fixed-point attractor is obtained with □ < 1.2, β = 3.6, α = 1.2, γ = 2.4, σ = 1.3 (Fig 2, panel b).
c. Sepsis progression and mortality Upon increasing LPS stimulation in either time span or amplitude, we assumed that *tlr4* mRNA rates are amplified proportionally, the result of which in our simulation leads to the system moving to a double-attractor. In this case TLR4 concentrations oscillate with highest amplitude and indefinitely between cell surface and EE compartments, with no signal resolution, using □ > 1.2, β = 3.6, α = 1.2, γ = 2.4, σ = 1.3 (Fig 2, panel c).

## 4. Discussion

Support for this overall scheme is found in emerging paradigms regarding sepsis progression and its lack of resolution. The overall immune response in sepsis is ultimately determined by a host of factors, chief among which are patient co-morbidities but crucially including also the virulence and size of the microbial inoculum. It has been proposed that following the initial LPS stimulation, the onset of sepsis in human patients encompasses both the pro-inflammatory and anti-inflammatory responses, with the former taking temporarily a more prominent role [34]. A composite cytokine score calculated to compare global inflammatory responses in murine models of sepsis demonstrated concomitant and similar upregulation of both pro-inflammatory and anti-inflammatory phases at 24 h before death [35]. We observed within the full course of this simulation asymmetrical oscillations residing preponderantly in a region of the phase space where TLR4 concentrations are augmented on the cell surface (site of pro-inflammatory signaling), before moving to a new region in phase space where similar variations are observed within the EE (with corresponding anti-inflammatory signaling), in a situation resembling the sepsis pathology. The outcomes for such fluctuations may lead to either death through the cytokine “storm”, or later on via the overall immunosuppression responsible for nosocomial infections and metabolic shutdown [36].

A post-hoc testing of this model using the initial, pre-treatment rates of *tlr4* mRNA from the patient cohort yielded appropriate descriptions of both the clinical outcome in 8 out of 10 patients, and the category of attractor each patient belongs to, as presented in Supplemental Table 1. Those patients whose TLR4 concentrations changes evolved towards one attractor were capable of surviving sepsis (patients #1, 4, 5, and 8). In contrast, those patients that presented a double-attractor state for TLR4 died within 3 days after ICU admission (patients #3, 6, 7, and 10). As a test to the sensitivity and specificity of our model, patient #2 died 9 days after ICU admittance due to *Candida albicans* infection. This pathogen is known to stimulate both TLR2 and TLR4, and is commonly associated with severe immunosuppression and high in-hospital mortality rates [37]. While TLR2 is an integral part to the initiation of the pro-inflammatory phase in sepsis and its blocking successfully rescues murine models from sepsis onset, we have not considered its role in this work. TLR2 and TLR4 co-stimulation with mycoplasma lipopeptides and LPS markedly increases tumor nuclear alpha production in macrophages, hallmark of synergy between these signaling pathways, thus complicating the use of TLR2 in our LPS-TLR4 only signaling model. Furthermore, patient #9 survived with negative microbiological cultures from both blood and pleural exudates. This may be the result of false-negative cultures, fungal infections, or the patient may have presented sepsis without the involvement of an infectious agent, a situation also not accounted for in our model.

## 5. Conclusions

We have used initial *tlr4* mRNA expression levels from sepsis patients in a dynamic model in order to describe the distribution of TLR4 within the cell surface compartment (pro‐inflammatory role), or intracellularly (anti-inflammatory and signal termination functions). We discriminated Gram-negative infections from the overall cohort and correctly predicted their clinical outcome. Confirming this hypothesis by in vivo measurements of TLR4 intracellular trafficking rates would provide further insight on their contribution to sepsis onset and progression.

## Author contributions

RSC analyzed the data and wrote the manuscript. FGS and MMDC prepared the experiments and performed data analysis. MMDC wrote and reviewed the manuscript.

## Conflict of interest

The authors report no conflicts of interest.

## Acknowledgements

We thank Dr. Grégoire Altan-Bonnet for critical reading of the manuscript. Financial support: CNPq (400662/2014-0 for RCS and 309041/2012-0 for MMDC), FAPESP (11/51778-6 for MMDC).

